# CrossModal Correspondence based MultisensoryIntegration: A pilot study showing how HAV cues can modulate the reaction time

**DOI:** 10.1101/2024.03.21.586134

**Authors:** Swati Banerjee, Daria Shumkova

## Abstract

We live in a multisensory world, where all our senses work together for giving us a fulfilling experience of the environment that we are in or during our use of immersive technologies.

For gaining more insight into the temporal scale understanding of the integration phenomenon EEG based BCI can give us the understanding of the transient changes in the brain.

In this study, we investigated the potential of incorporating haptics into crossmodal correspondence based research to induce MSI effect through either the active touch users’ feedback or crossmodal correspondences with visual and auditory modalities, such as Kiki Bouba effect.

We designed two experiments:

1. Visual stimuli were presented on a standard computer monitor, and auditory stimuli were delivered through computer dynamics. Participants responded using left or right hand by pressing either CapsLock or Enter buttons respectively. Visual cue consisted of a red circle displayed randomly either on the left or on the right side of the screen. Auditory cue was a brief high tone presented through left or right headphones for 500 ms. Text stimuli that appeared on the screen instructed participants to respond with their left or right hand. Before each trial there was a fixation central cross displayed for 500 ms.
2. This experiment was inspired by previous studies on Kiki-Bouba correspondence. Visual stimuli consisted of 4 shapes - circle, triangle, polygon with 6 vertices, and star - presented on a computer screen. Locations of the visual stimuli were randomized. Auditory stimuli were generated using the Online Tone Generator website (https://onlinetonegenerator.com/). 2 sets of sounds were used: the first set included sine, triangle, square, and sawtooth waveforms, each at a frequency of 500 Hz; the second set included sawtooth waveforms at frequencies of 50 Hz, 300 Hz, 600 Hz, and 2000 Hz (summarised in Table 2).

Results suggested that it is indeed possible to achieve this type of integration without relying on complex haptic devices. Introducing haptics into BCI technologies through feedback touch or crossmodal correspondances holds potential to improve the user experience and information transfer rate (ITR).

Participants, as expected, showed the lowest reaction times in congruent sequential test and the highest – in incongruent HAV cues based test. This indicates the importance preference for sequential cue presentation over simultaneous one. The time was significantly higher in case of Incongruent Haptic cues.

## 1 Introduction

We live in a multisensory world, where all our senses work together for giving us a fulfilling experience of the environment that we are in or during our use of immersive technologies. Signals from various sensory pathways are co-processed and integrated to enhance neuronal activity, leading to MultiSensory Integration (MSI) [1]. On the physiological level, MSI can happen in the brain areas receiving afferent inputs from several sensory pathways. The effect of MSI is not limited to these specific brain regions. It was shown that MSI can also happen in the single sensory brain regions and affect various brain functions [1]. Previous studies have shown that MSI starts after 50 ms, after 46 ms and after 80 ms of the stimulus onset for the auditory-tactile, auditory-visual and visual-tactile stimuli respectively [2]. Moreover, Talsma et al. reported that attention can influence MSI at the different stages of processing [3]. Technological advancements in the last decades have introduced new modalities in Brain-Computer Interfaces (BCIs). Despite this, many BCI based applications still predominantly uses single modality in terms of Cues presentations in case of technology integration. Specifically visual stimuli still remains irreplaceable as human tend to react to respond to visual representation better than any other form of stimuli. For gaining more insight into the temporal scale understanding of the integration phenomenon EEG based BCI can give us the understanding of the transient changes occuring in the brain. The main idea behind this type of study and this work is to improve the performance and also to create more efficient and ecological protocols, where we aim to design stimuli resembling the real world sensory integration. These stimuli are basically cues arriving in form of signals that can potentially improve the integration of senses. Within the scope of this study we specifically focused on the reaction time and a potential improvement of the same in terms of accuracy, this can give us a better understanding of the coding decoding phenomenon going on in the brain micro states. One widely know and explored way of achieving the above mentioned task is Cross Modal Correspondence (CMC) which is briefly explained below.

### Cross Modal Correspondences

One way how brain can facilitate MSI is through Cross Modal Correspondence. These exists stable associations of stimuli from the different sensory modalities that we tend to share with the other people as a “common ground” [4] for joint actions in case of group activities in terms of shared information that provides the required basis for mutual understanding and performing a task. For instance, individuals tend to link high-pitched sounds with small, light objects and low-pitched sounds with large, dark objects. An example of such association made between the stimuli in the auditory and visual modalities is given in Fig. 1. Another example of crossmodal correspondances is the link between shape of the food and its taste [5]. In the context of somatosensory modality, temperature was shown to be associated with colour [6]. In the recent study, Kanaya et al., found touch-vision and touch-audition correspondances [7]. Together it shows that crossmodal associations are possible between any kind of sense combinations. Regarding the function of this phenomenon, it is hypothesised that these shared associations assist in choosing which senses should be processed together as the part of the MSI [4]. More practically speaking, if the BCI stimuli in one’s study are linked through CMS correspondance, we hypothesise that the neuronal response of the BCI user can be improved through MSI.

**Figure 1.**
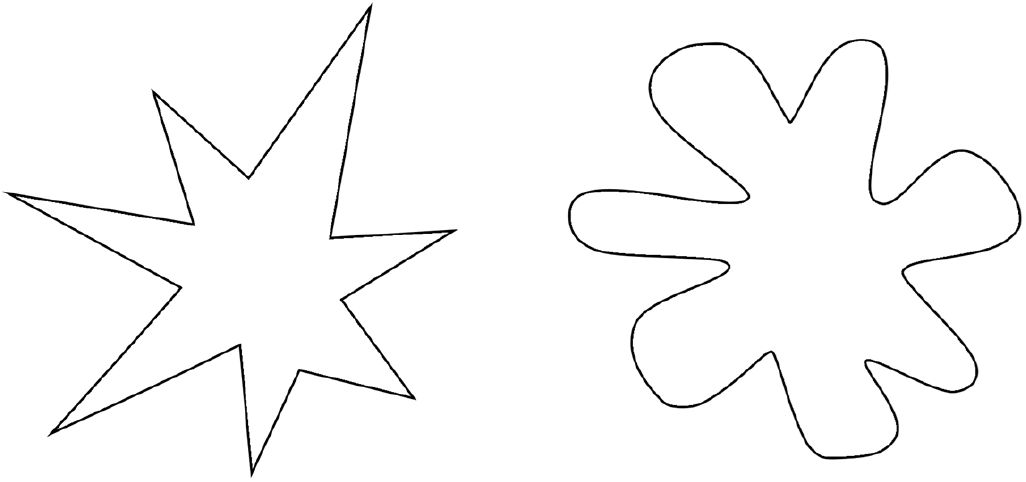
Kiki-Bouba effect. People tend to associate the left shape with the high tones and the right shape with the low tones.

### MultiSensory Integration in Brain Computer Interfaces

EEG-based BCIs were designed to allow e.g. paralysed patients to spell out words by selecting letters on a virtual keyboard using their brain’s responses. There are several types of brain responses that can be used by decoders. The most widely studied P300 potential occurs at 300 ms after stimulus demonstration and is usually a positive deflection of voltage in the EEG trace. Steady-State Evoked Potentials (SSEP) is another brain pattern that appears upon the stimulus presentation and can be used in BCIs as an alternative to P300 to increase its efficacy [8]. Unlike P300 SSEP are relatively constant and periodic, occuring on the duration of longer time period. Moreover, SSEP is evoked by continuous or repetitive stimulation, while P300 is elicited even after a single short stimulus. The main rationale of developing the multimodal BCIs is to check whether it can outperform visual-only BCIs which are not suitable for the use of many patients e.g. with disorders of visual system. Table 1 provides an overview of the crossmodal EEG studies showing how multimodal approaches influence the BCI performance. We have taken into consideration all this previous work to as a basis of our stimuli design experiment. It is still very difficult to present

**Table 1:** Stimuli presented in sequential or simultaneous manner across different studies on crossmodal EEG. V - visual, A - auditory, H - haptic/tactile

| Study | Modality used | P300 | Other evoked potentials |
| --- | --- | --- | --- |
| Thurlings et al., 2012 | Bimodal HV | Unimodal V > Bimodal HV | Enhanced bimodal HV N1 |
| Thurlings et al., 2014 | Bimodal HV | Bimodal HV > Unimodal V | - |
| Brouwer et al., 2010 | Bimodal HV | Bimodal HV > Unimodal V | - |
| Yin et al., 2016 | Bimodal HA | Bimodal HA > Unimodal | - |
| Belitski et al., 2011 | Bimodal AV | Bimodal AV > Unimodal | - |
| Lu et al., 2019 | Bimodal AV | Bimodal AV > Unimodal | Enhanced bimodal AV P600 |

### Adding Haptics based sensations in Brain-Computer Interfaces

Auditory and visual cue modalities have long been an integral components of various technologies and studied particularly in BCI research. However, the incorporation of haptic feedback has only recently captured the attention of researchers with the increase in the haptic and tactile based emergent technologies. This type of haptic based paradigms always engages the somatosensory pathway. In the context of motor imagery tasks, the closure of the sensorimotor loop through haptic feedback holds significant promise for motor rehabilitation applications by promoting plasticity mechanisms [8]. Additionally, there is a growing suggestion that integrating haptic feedback in BCIs could enhance the subject’s sense of agency, a crucial factor in technology acceptance. Despite the potential benefits, haptic feedback remains underutilized in the BCI/neurofeedback technologies, even though the haptic sense uniquely enables us to interact with and perceive the world around us. The challenge lies in the complexity of integrating haptics into real-world BCI applications, as it demands sophisticated devices. Moreover, some vibrotactile devices, commonly used for haptic feedback, introduce mechanical waves that generate unintended noise, posing a challenge in experimental contexts. These technological constraints also make it difficult to ensure full congruence between haptic stimuli and other sensory inputs, limiting the seamless integration of haptics into BCIs. In this project we hypothesised that haptics or haptics like sensations can be introduced along with the visual and auditory modalities without the sophisticated specialised devices through low-tech solutions.

To check this hypothesis two independent MSI experiments were designed. In the experiment 1 haptics was introduced through active users’ feedback touch of the keyboard with either left or right hand in response to the other stimuli.

In the experiment 2 Kiki-Bouba crossmodal correspondence was reproduced, where haptics is thought to be linking the auditory and visual modalities (e.g. through shape imagery [9]). We introduced the active touch component of the study by keystroke events.

## 2 Methods

Both experiments were designed using PsychoPy software (Version 2023.2.3; [10]). Demographic information, such as age, gender, and any relevant background characteristics, was recorded.

For each of the experiments, trial-level data were recorded, including reaction times - the time difference between the onset of the last stimulus and the participant’s response; and the accuracy data, indicating whether the participant’s response was correct or incorrect. For the second experiment the recorded data also included the matched visual-auditory pair. Statistical analysis for this project included the unpaired Student’s t-test, with significance set at a threshold of p *<* 0.05.

### Experiment 1: Feedback Touch

Visual stimuli were presented on a standard computer monitor, and auditory stimuli were delivered through computer dynamics. Participants responded using left or right hand by pressing either CapsLock or Enter buttons respectively. Visual cue consisted of a red circle displayed randomly either on the left or on the right side of the screen. Auditory cue was a brief high tone presented through left or right headphones for 500 ms. Text stimuli that appeared on the screen instructed participants to respond with their left or right hand. Before each trial there was a fixation central cross displayed for 500 ms.

Training consisted of 30 trials, evenly distributed between 3 phases (see Fig. 2A) Visual Training (10 trials): Participants responded to the location of a red circle by pressing the corresponding left or right button. Auditory Training (10 trials): Participants responded to the side of the headphone from which an auditory cue was given. Haptics Training (10 trials): Participants responded according to the instruction presented on the screen.

**Figure 2.**
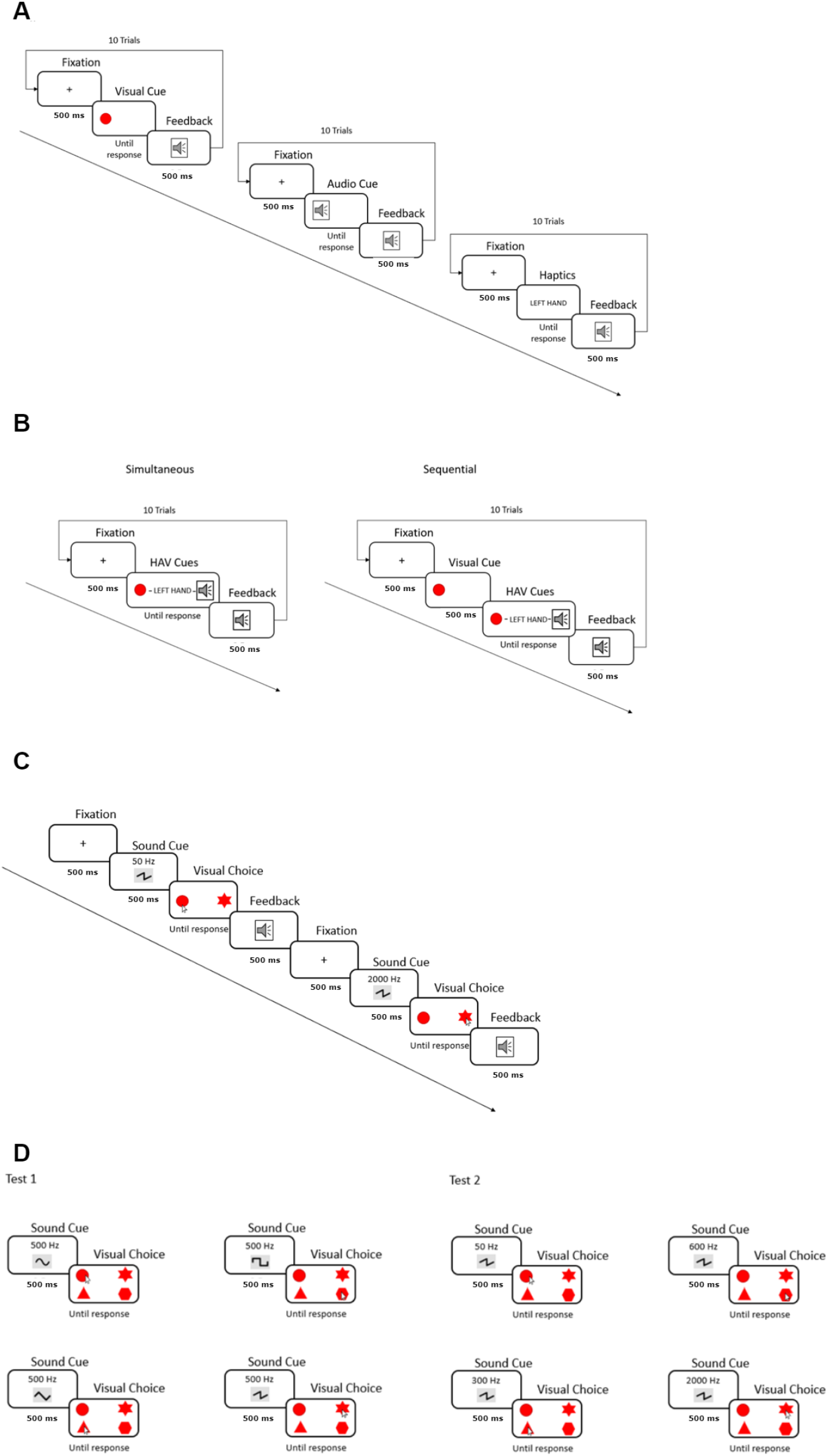
Experimental pipeline for training (A, C) and testing (B, D) settings. In the experiment 1 (A,B) haptics was introduced through keyboard touch in response to text-stimulus, visual cue was a red circle target and auditory cue was a brief high tone. In experiment 2 (C,D) Kiki-Bouba crossmodal correspondence was introduced through forced matching of the shapes of different angularity and tones of different frequencies/waveforms.

Testing consisted of 40 trials (see Fig. 2B). Congruent Trials (20 trials): Visual, auditory, and text cues were presented simultaneously (10 trials) or sequentially (10 trials where auditory cue and response instruction were 500 ms delayed relatively to visual cue) in a congruent manner (e.g. left visual cue, left headphone tone, left hand response), and participants responded accordingly. Incongruent Auditory Trials (10 trials): Auditory cues were incongruent with visual and text cues, and participants were instructed to respond based on the visual and text information. Incongruent Hand Response Trials (10 trials): Participants were instructed to respond with the hand opposite to the information provided by visual and auditory cues.

### Experiment 2: Crossmodal Correspondance

This experiment was inspired by previous studies on Kiki-Bouba correspondence. Visual stimuli consisted of 4 shapes - circle, triangle, polygon with 6 vertices, and star - presented on a computer screen. Locations of the visual stimuli were randomized. Auditory stimuli were generated using the Online Tone Generator website (https://onlinetonegenerator.com/). 2 sets of sounds were used: the first set included sine, triangle, square, and sawtooth waveforms, each at a frequency of 500 Hz; the second set included sawtooth waveforms at frequencies of 50 Hz, 300 Hz, 600 Hz, and 2000 Hz (summarised in Table 2).

**Table 2:** Visual and auditory stimuli used in Experiment 2

|  | Visual Shape | Matched Sound frequency, Hz | Matched Sound waveform |
| --- | --- | --- | --- |
| Training | Circle<br>Star | 50<br>2000 | Sawtooth |
| Test 1 | Circle<br>Triangle<br>Polygon<br>Star | 500 | Sine<br>Triangle<br>Square<br>Sawtooth |
| Test 2 | Circle<br>Triangle<br>Polygon<br>Star | 50<br>300<br>600<br>2000 | Sawtooth |

Participants began with a training session to familiarize themselves with the visual shapes and the corresponding sounds (see Fig. 2C). 12 trials were conducted with visual stimuli (circle and star) paired with auditory stimuli (50 Hz and 2000 Hz sawtooth tones). Tones’ distribution through trials was randomised.

The testing comprised 2 phases: in test 1 - the sound frequencies were altered; in the test 2 - the sound waveforms were modified, with the frequency remaining constant (see Fig. 2D). In the first test, participants completed 24 trials where they were presented with the 4 visual shapes (circle, triangle, polygon with 6 vertices, and star) and were asked to match each shape with 1 of 4 sounds, all at a frequency of 500 Hz. The sounds had different waveforms: sine, triangle, square, and sawtooth. First 12 trials were non-random, where the sounds were played in the following order: sine, triangle, square and sawtooth waveforms. In the last 12 trials sounds were randomized.

The second test involved another set of 24 trials. Participants were presented with the same 4 visual shapes and were required to match each shape with one of 4 sawtooth waveforms, each at a different frequency (50 Hz, 300 Hz, 600 Hz, and 2000 Hz). First 12 trials were non-random, where the sounds were played in the ascending frequency order. The last 12 trials were randomized.

## 3 Results

### Participants

12 subjects participated in the experiment 1 with the mean age of 25.2 (±2.0): 9 males and 3 females. In the second experiment 16 subjects were recruited with the mean age of 27.7 (±8.1): 9 males and 7 females (see Table 3). All participants were right-handed and did not have any visual or auditory complications. It is important to note that the participants were different between the 2 experiments.

**Table 3:**
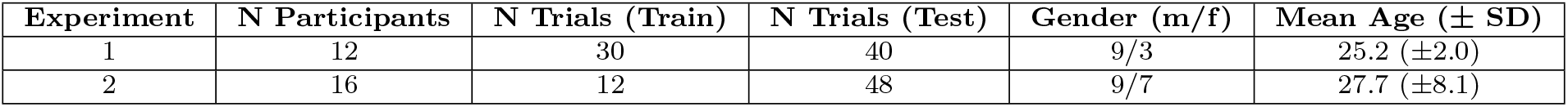
Demographic information about participants

### Experiment 1. Incongruency of haptic feedback caused suboptimal performance in MI tasks

From Fig. 3 it becomes evident that participants in general improved their performance throughout the trials for each of the tasks in the experiment 1. Learning curves in congruent simultaneous stimuli presentation showed a relatively rapid improvement over time compared to all the incongruent tasks. Incongruent and congruent sequential tasks followed 30 training trials and 10 congruent simultaneous trials, meaning that the users were already familiar with the job and the instructions were clear. It was still observed that performance in incongruent tasks was suboptimal, particularly when the users were required to provide feedback using the opposite hand.

**Figure 3.**
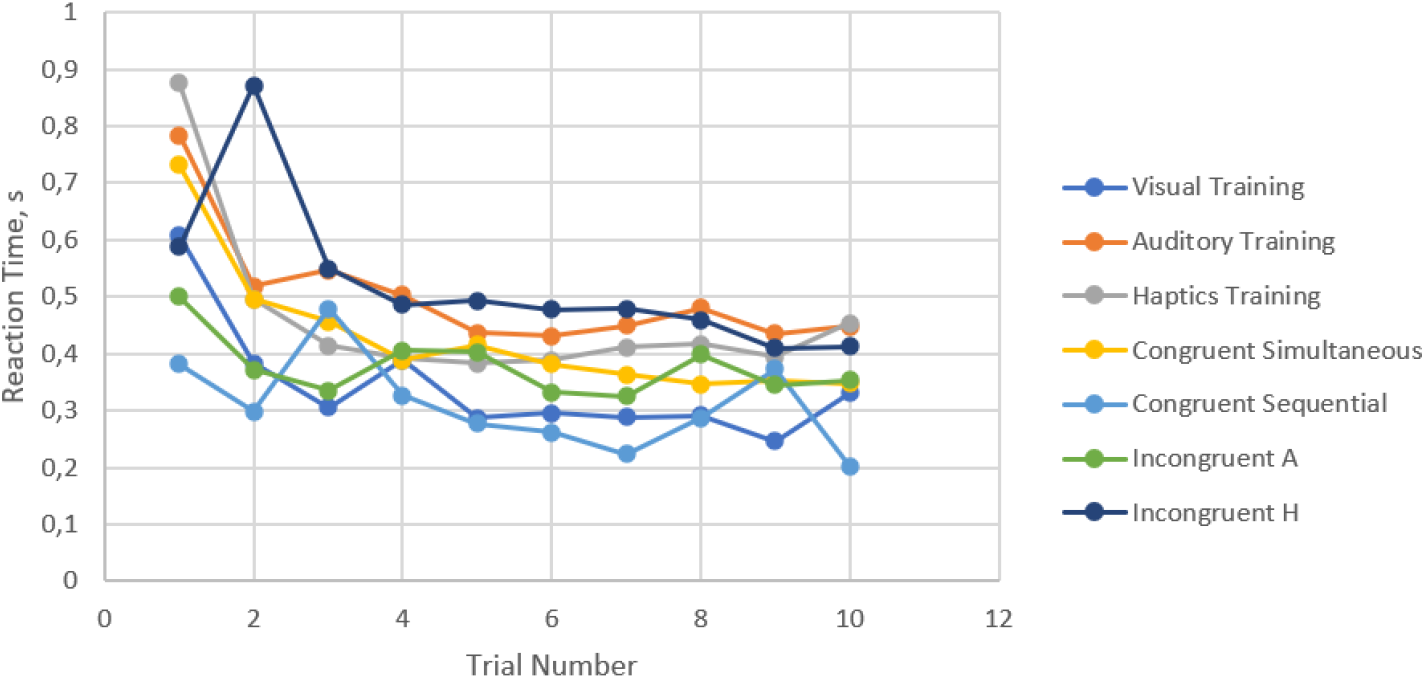
Learning curve for the experiment 1 across trials averaged between participants. A - auditory, H - haptics.

Moreover, in line with the learning curve results, the lowest accuracy and the highest mean reaction times were observed in the incongruent haptics task (see Fig. 4). Together this indicated that congruency of haptic cues is vital to ensure the optimal perfomance of the subjects in the MI tasks.

**Figure 4.**
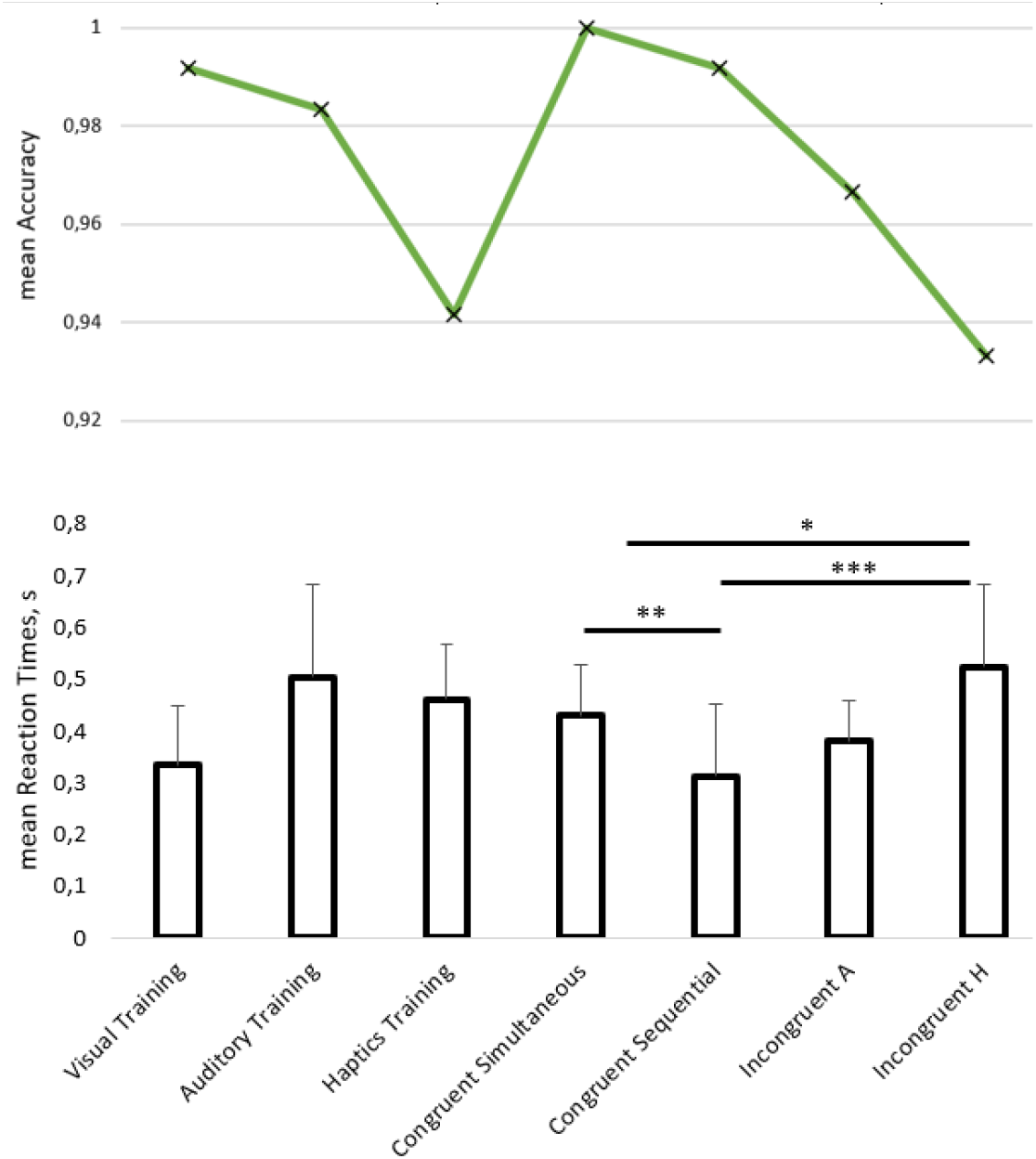
Mean reaction times and accuracies in experiment 1 across different tasks, ^*^p¡0.05, ^**^p¡0.01,^***^p¡0.001 in paired t-test. Outliers were excluded from the reaction times. A - auditory, H - haptics.

### Experiment 2. Stimuli matched through crossmodal correspondances seem to induce Multisensory Integration

The data structure for experiment 2 is summarised in Fig. 5A. In the second experiment learning curve indicates that participants notably improved their performance over trials only in the non-random test 1 setting, where auditory stimuli were presented in the order of increasing sharpness of their waveforms (see Fig. 5B). In the other testing conditions subjects’ performance was fluctuating, meaning they did not develop a clear strategy of shape-sound matching and their choices seemed “random” to themselves.

**Figure 5.**
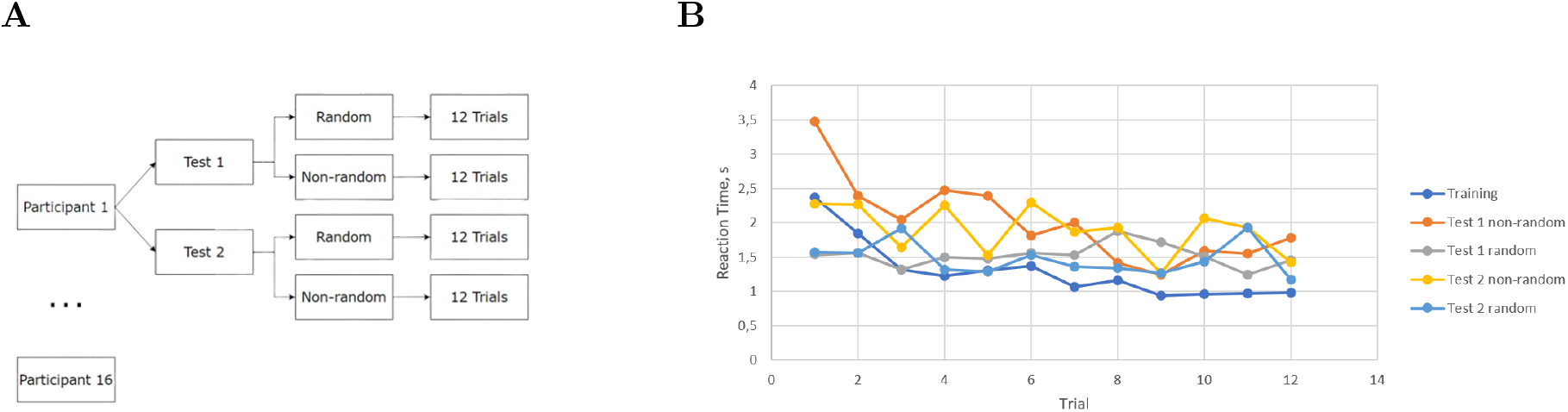
Data structure of the experiment (A) and learning curve for experiment 2 (B) across trials averaged between participants.

**Figure 6.**
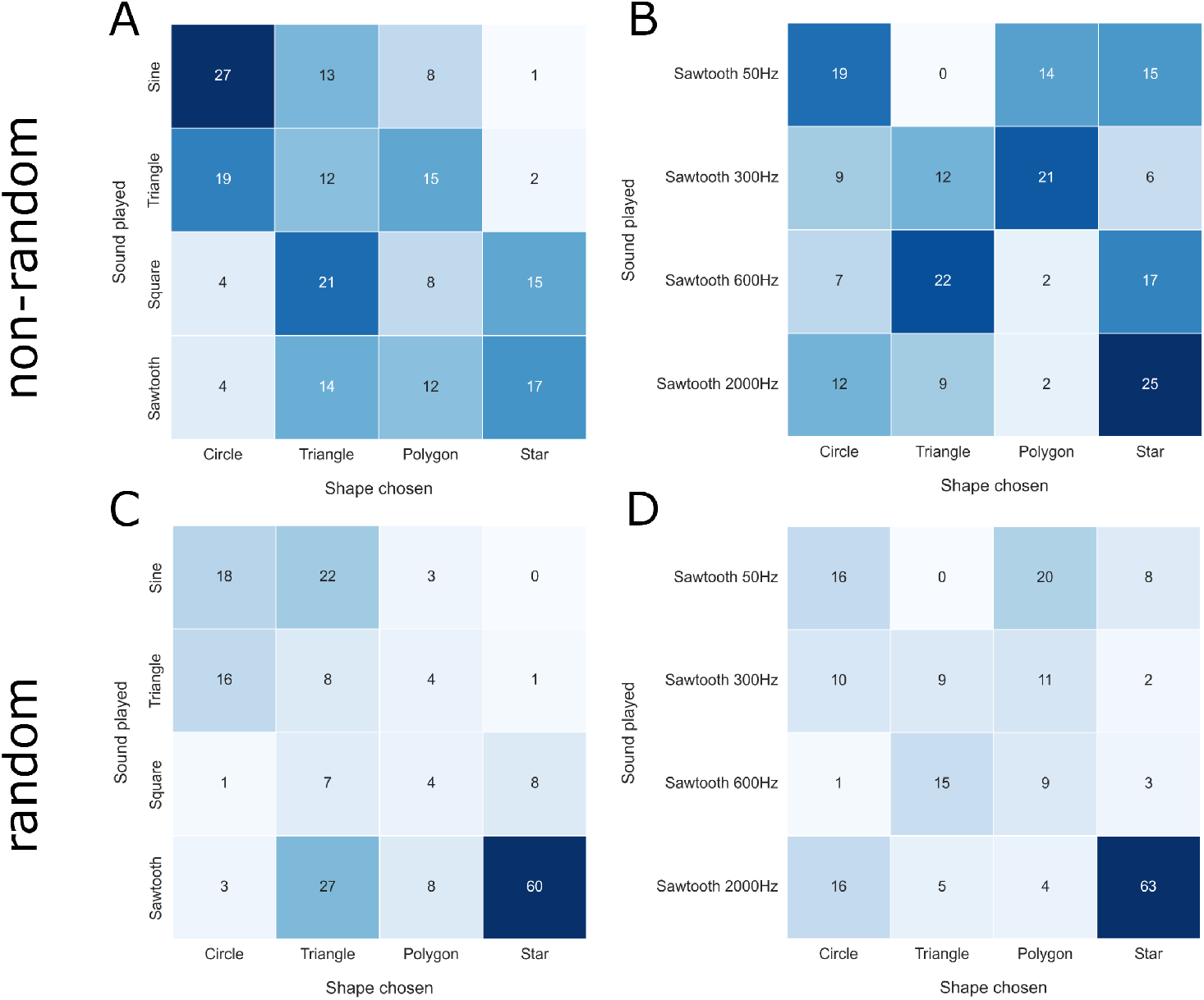
Confusion matrices showing the total number of matches between different shapes and sounds made by the participants. Auditory stimuli were presented either (A,B) non-randomly across trials or (C,D) in a completely random fashion. Moreover, sounds were different either by (A,C) waveform or (B,D) frequencies.

From the confusion matrices it becomes evident that star shape was the easiest to be matched regardless of the testing condition (see Fig. 7). As expected, sharper sound waveforms (e.g. 2000 Hz, sawtooth) were matched to sharper targets (e.g. triangle, star), indicating the effect of Kiki-Bouba crossmodal correspondence. In general participants performed better when the auditory stimuli were varying by the waveforms rather than the frequencies. In part it can be explained by a poor choice of 50 Hz auditory stimulus that might have caused unintended buzzling sensation in participants.

## 4 Discussion and Conclusion

In this study, we investigated the potential of incorporating haptics into research to induce MI effect through either the active touch users’ feedback or crossmodal correspondences with visual and auditory modalities, such as Kiki Bouba effect. Results suggested that it is indeed possible to achieve this integration without relying on complex haptic devices. Introducing haptics into BCI technologies through feedback touch or crossmodal correspondances holds potential to improve the user experience and information transfer rate (ITR).

